# Regional activity in the rat anterior cingulate cortex and insula during persistence and quitting in a physical-effort task

**DOI:** 10.1101/2020.05.25.115576

**Authors:** Blake S. Porter, Kunling Li, Kristin L. Hillman

## Abstract

As animals carry out behaviors, particularly costly ones, they must constantly assess whether or not to persist in the behavior or quit. The anterior cingulate cortex (ACC) has been shown to assess the value of behaviors and to be especially sensitive to physical effort costs. Complimentary to these functions, the insula is thought to represent the internal state of the animal including factors such as hunger, thirst, and fatigue. Utilizing a novel weight lifting task for rats, we characterized the local field potential (LFP) activity of the ACC and anterior insula (AI) during effort expenditure. In the task male rats are challenged to work for sucrose reward, which costs progressively more effort over time to obtain. Rats are able to quit the task at any point. We found modest shifts in LFP theta (7-9 Hz) activity as the task got progressively more difficult in terms of absolute effort expenditure. However, when the LFP data were analyzed based on the rat’s relative progress towards quitting the task, or performance state, substantial shifts in LFP power in the theta and gamma (55-100 Hz) frequency bands were observed in ACC and AI. Both ACC and AI theta power decreased as the rats got closer to quitting, while ACC and AI gamma power increased. Furthermore, coherency between ACC and AI in the delta (2-4 Hz) range shifted alongside the rat’s performance state. Overall we show that ACC and AI LFP activity changes correlate to the rats’ relative performance state in an effort-based task.

**Significance Statement:** Animals need to assess whether or not a behavior is worth pursuing based on their internal states (e.g., hunger, fatigue) and the costs and benefits of the behavior. However, internal states often change as behaviors are carried out, such as becoming fatigued, necessitating constant reassessment as to whether to continue the behavior or quit. We characterized brain activity in the anterior cingulate cortex and insula, brain regions involved in cost-benefit decision making and internal state representations, respectively, as rats carried out a challenging physical-effort task. Both brain regions showed significant shifts in activity as the rats approached their quitting point. Our study provides one of the first characterizations of neural activity as an animal decides to quit an effortful task.

## Introduction

When tasks become effortful, animals must evaluate whether additional physical and/or mental effort expenditures are worthwhile. This evaluation requires both a cost-benefit assessment of the task and an interoceptive assessment – an animal must gauge physiological and/or psychological preparedness to handle any continued effort expenditure. While multiple brain regions are involved in these processes, two regions appear central: the anterior cingulate cortex (ACC) and the anterior insula (AI).

Based on lesion and electrophysiological studies, the ACC is implicated in effort-based cost-benefit analysis. For example, when rats are presented behavioral options that include varying levels of effort, ACC single unit activity is reflective of the behavior with highest utility (Hillman and Bilkey 2010; Cowen et al. 2012; Hart et al. 2019; Porter et al. 2019). Human EEG recordings indicate that frontal midline theta, thought to originate from the ACC, correlates to task effort and difficulty (Smit et al. 2005; Cavanagh and Frank 2014). Taken together, heightened population activity in the ACC may help drive action towards high-utility goals when effort is required (Amiez et al. 2006; Kennerley and Wallis 2009; Cowen et al. 2012; Hillman and Bilkey 2012).

The AI is implicated in representing the interoceptive state of the body. The insula receives hypothalamic, autonomic, and visceral inputs making the AI well-positioned to process interoceptive information on states such as hunger and pain, which can influence cost-benefit decision-making (Cechetto and Saper 1987; Craig 2003; Critchley 2005). In rats, lesions to the AI result in maladaptive persistence and perseveration behaviors in reward devaluation scenarios (Balleine and Dickinson 2000; Parkes et al. 2015; Moschak et al. 2018). One interpretation of this is that AI functionality is needed during ongoing cost-benefit decision making to spur effort reallocation – in the form of task switching and/or quitting – when a behavior has decreased utility relative to the body’s needs. In turn, the AI’s interoceptive state representation could be utilized by the ACC to evaluate the costs and benefits of persisting at or quitting a behavior.

White matter tracts between the cingulate and insula have been traced in humans (Moisset et al. 2010), and resting state functional connectivity between the regions have been reported (Sridharan et al. 2008; Taylor et al. 2009). As outlined by Medford and Critchley (2010) the regions could function in tandem: the AI integrating external and internal sensory inputs to represent the current state of the organism, with the ACC then utilizing this state representation to drive relevant behaviors. If external and internal sensory inputs to the AI increasingly signal effort costs (fatigue, energy depletion), the ACC could bias behaviors away from continued energy expenditure. Both brain regions have been shown to encode the expected energetic cost of physically demanding behaviors (Prevost et al. 2010), and as tasks get increasingly effortful the ACC and AI show conjoint BOLD activation (Engstrom et al. 2014). It is not well-understood, however, how dynamic changes in conjoint ACC-AI activity correlate to changes in effortful task performance. Furthermore, few studies have examined ACC-AI activity to the point of quitting.

To address these questions we recorded local field potentials (LFPs) from the ACC and AI of rats as they performed a novel weight lifting task (WLT). The WLT allows systematic manipulation of physical effort costs while maintaining the same motor pattern, spatial context, and reward. LFPs were analyzed for shifts in power across the major frequency bands: delta (2-4 Hz), theta (7-9 Hz), beta (15-25 Hz), and gamma (55-100 Hz). Oscillation frequency is thought to reflect the number of synchronous neurons engaged in a given process – higher frequencies reflect local activity while slower rhythms reflect the synchronous activity of neurons across brain regions (von Stein and Sarnthein 2000; Buzsaki and Draguhn 2004). We predicted that the ACC and AI may be communicating via low-frequency oscillation during an effortful task, and that low-frequency LFP power from the ACC would inversely correlate to effort costs to reflect the decreasing utility signal generated by the ACC. In contrast, low-frequency LFP power from the AI would positively correlate to effort costs as fatigue increases and internal states drive behavior away from task engagement. In addition, we investigated the potential involvement of the beta rhythm, often associated with sensorimotor systems (Salenius and Hari 2003) and working memory (Engel and Fries 2010), and the gamma rhythm, thought to reflect local circuit activity (Buzsaki and Draguhn 2004; Bartos et al. 2007; Merker 2016). We also characterized ACC-AI LFP activity at the quitting point, a time period that to our knowledge has been minimally investigated at the electrophysiological level.

## Materials and Methods

### Subjects

Male Sprague-Dawley rats (n=16, weighing 400 - 575 g at the time of experiment; Hercus Taieri Resource Unit, New Zealand) were single housed in individually ventilated cages, 38 × 30 × 35 cm (Tecniplast, Italy). Animals were maintained on a 12 h reverse light cycle, with experimentation occurring during the animals’ dark, active phase. Water was available *ad libitum*. Daily chow (Teklad diet; Envigo, USA) was rationed to promote interest in food reward during experimentation. Rats were weighed twice per week and daily food rations were adjusted to maintain each animal’s body weight ≥ 85% of their free-feeding body weight. All procedures were approved by the University of Otago’s Animal Ethics Committee, protocol 91/17.

### Training

Prior to surgery, animals were trained in the WLT as previously described (Porter and Hillman 2019). Briefly, the WLT occurs in a 120 × 90 × 60 cm arena (Figure 1a & 1b); an animal is progressively shaped to pull a rope 30 cm to trigger dispensing of 0.25 mL of 20% sucrose solution. The rope can be weighted from 0 to 225 g to increase the effort demands of the task, however only 0 g and 45 g are used in the training phase. Once an animal can perform 20 successful pulls (10 pulls on 0 g followed immediately by 10 pulls on 45 g) in under five min, they are considered trained in the WLT and ready for surgery.

**Figure 1:**
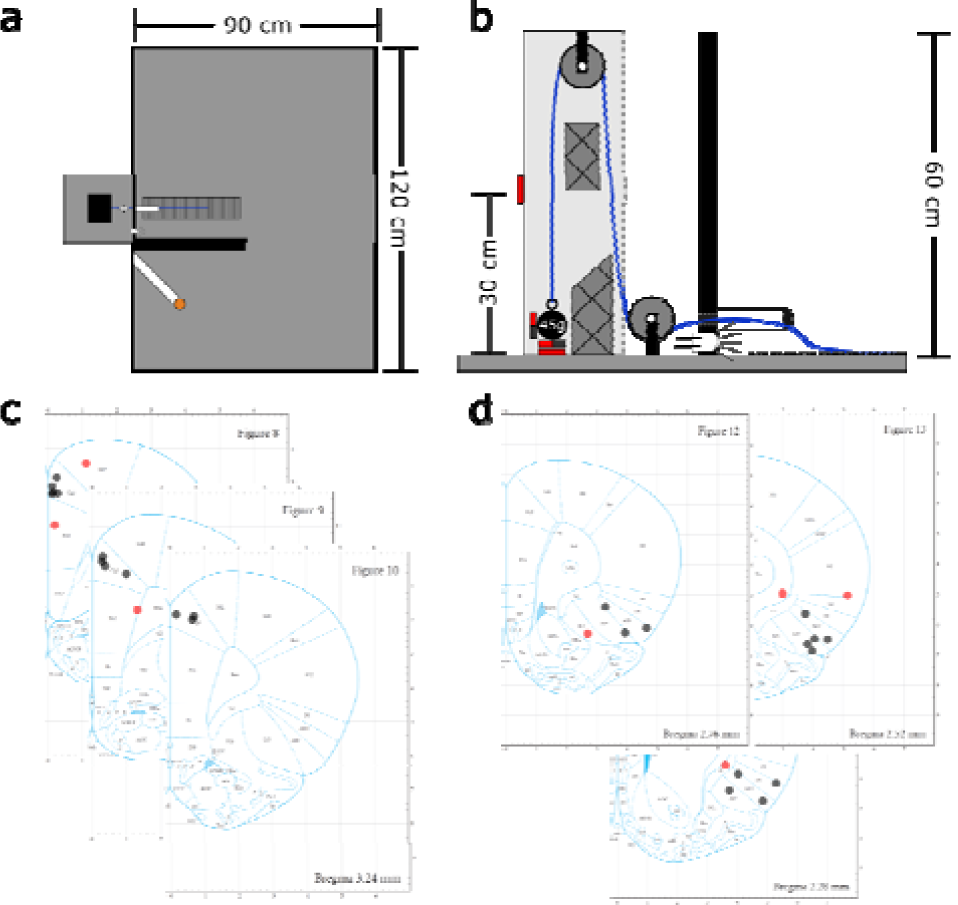
Experimental overview. a) Weight lifting task apparatus. The pulley system is depicted by the black box on the left, outside the arena. The rope is shown in blue extending into the arena over the rubber grip pad (vertical lines). The 20% sucrose reward is delivered to the orange cap separated from the rope by the black barrier. b) The weight lifting pulley schematic. The rope is shown in blue with one end attached to a 45 g weight with two magnets and the other end extending into the arena through the conduit. The hash mark shapes indicate Styrofoam inserts to prevent the weight from swinging while pulled. The magnetic sensors are shown in red. c-d) Histological reconstructions illustrating electrode tip locations for the ACC (c) and AI (d). Black dots were on-target placements while off-target placements are red. Off-target electrodes were not used for LFP analyses. Panels (a-b) adopted from Porter and Hillman (2019); panels (c-d) adopted from Paxinos and Watson (2007).

### Surgery

Animals were implanted with two, 200 µm insulated nichrome electrodes (Johnson Matthey PLC, UK) under isoflurane anesthesia as previously described (Porter et al. 2019). Stereotaxic coordinates, based on Paxinos and Watson (2007), were set as follows: For the ACC we targeted Cingulate Cortex area 1 (Cg1; equivalent to region 32d) at + 3.7 mm anteroposterior (AP) from Bregma, + 0.4 mm mediolateral (ML), - 1.0 mm dorsoventral (DV); and AI + 2.7 mm AP from Bregma, + 2.0 mm ML, and - 5.8 mm DV at an angle of 20° ML. Dorsoventral depths are from brain surface. All implants were in the right hemisphere. A 4 mm stainless steel screw placed over the cerebellum served as the ground connection. Five to seven additional 3 mm stainless steel screws were placed in the skull for dental cement to anchor to. Electrodes were secured with UV cure self-etching dental cement (Maxcem Elite™, Kerr Corp., USA) followed by additional self-cure dental acrylic (Lang Dental Manufacturing Company, Inc., USA) for additional strength. Ten days after surgery and post-operative care, rats were retested on the WLT with 0 and 45 g to ensure no loss of function from surgery and to check the quality of the electrophysiological signals. Within 3 days, all rats returned to baseline performance criterion (i.e., 10 successful trials on 0 g and 10 successful trials on 45 g, all within 5 min). Once criterion performance was confirmed, experimentation began.

### Weight Lifting Task

To test persistent, effortful behavior, a progressive weight lifting paradigm was utilized. This paradigm has been detailed previously (Porter and Hillman 2019). Briefly, the rats need to pull a rope with a weight attached for 30 cm to trigger release of 0.25 mL of 20% sucrose solution. In the progressive WLT, an animal is challenged to pull progressively heavier weights: the weight starts at 0 g and after every 10 successful, rewarded pulls, 45 g of weight is added to the rope. The animal continues until they quit or a rope weight of 225 g is reached. If 10 successful pulls are achieved on 225 g then the weight is made ‘impossible’ by affixing it to the set-up such that the weight can only be pulled up approximately 15 cm, thus making it impossible to trigger a reward. Quitting was defined as 2 minutes of no attempted rope pulls.

Each task session was bookended with 2 min of open field exploration in the arena. During these open field periods the rope was not available and no sucrose was dispensed. The initial 2 min exploration provided a baseline measure of LFP and locomotor activity prior to beginning the WLT. Likewise, the 2 min exploration at the end of the task provided post-task LFP and locomotor measurements. At the end of the 2 min post-task exploration, a sucrose reward check was performed to test for satiation. The sucrose satiation check entailed two manually released sucrose rewards and the rats had one minute to approach and consume the sucrose. Whether or not they approached and consumed the sucrose was noted by the experimenter.

### Recordings

Local field potentials were recorded using a Neuralynx (USA) acquisition system (either a Digital Lynx SX or SX-M) at 6,400 or 5,000 Hz, respectively, with a bandpass filter set between 0.1 and 500 Hz. Rats were tethered via a Neuralynx Saturn-1 commutator to either a Neuralynx HS-36-LED or HS-36-mux-LED pre-amp headstage. Rat position was tracked via two headstage mounted LEDs by Neuralynx’s Cheetah software using an overhead camera. Rats were ran in a darkened room to allow tracking of the LEDs. The Neuralynx acquisition system also recorded TTL signals from the WLT Arduino Uno microcontroller (Arduino.cc) that timestamped when the weight was lifted off the base, when the weight returned down to the base, and when the weight was successfully pulled up 30 cm and a reward was released.

### Histology

At the end of the study, animals were transcardially perfused with 4% paraformaldehyde in PBS and histology performed as previously described (Porter et al. 2019). Thionin staining confirmed 12 out of 16 correct placements in the ACC (Figure 1c), and 12 out of 16 in the AI (Figure 1d). Every rat included for analyses (behavioral and electrophysiological) had at least one electrode in one of the target regions (ACC or AI). Of those excluded: one ACC electrode could not be confirmed due to the tissue being damaged during histological preparation; three ACC targeted electrodes were confirmed to be misplaced; four AI targeted electrodes were confirmed to have missed the AI. LFP data from electrodes with incorrect placements were excluded from analysis. Behavioral data was analyzed for all 16 rats. Note there is continued debate on the rodent equivalent of the primate ACC, see Laubach et al. (2018) for review, and our ACC electrodes are primarily in ACC area 1 / 32d.

### Experimental Design and Statistical Analysis

Statistical analysis was carried out in MATLAB with custom and native scripts. All statistical analyses were double checked in Prism (Version 8.4.1; GraphPad Software). Behavioral and LFP data were parsed based on the timestamps relayed by the Arduino TTL signals and experimenter notes using MATLAB. Methods for behavioral analyses have been reported in detail previously (Porter and Hillman 2019). Briefly, data from a given behavioral metric was first tested for normality using a one sample Kolmogorov-Smirnov test. Depending on the data’s distribution the appropriate statistical test was selected, either a one-way ANOVA or Kruskal-Wallis test, with a P value of 0.05. Dunn-Sidak’s Method was used for making post-hoc multiple comparisons.

### LFP analyses

Each rat (n=16) ran the progressive WLT for 5 to 8 recording sessions, one session per day. In total there were 83 recording sessions for ACC data and 82 for AI. Each recording session was broken up into epochs based on the different rope weights (0, 45, 90 g, etc.). Each epoch began when the respective weight was attached to the rope, and ended when the next weight was attached after 10 successful trials or until the rat quit. The 0 g epoch began immediately following the 2 min pre-baseline period when the rope was extended into the apparatus. Our LFP analyses focused on rope pull attempts. LFP attempt epochs were defined as the 2 sec preceding the rat lifting the weight, i.e., the initiation of task execution. If attempts overlapped in series (e.g., the rat pulled the weight, dropped it, and then pulled again within 2 sec) only the first attempt was used. Attempts were analyzed regardless of their outcomes (successful or failed pull). If the rat did not have at least five attempts for a given weight, that weight was not included in the LFP analysis.

LFP signals were down sampled to 200 Hz prior to analysis using Matlab’s downsample function. LFP data were then analyzed using the Chronux toolbox (Bokil et al. 2010). After down sampling, LFP signals were locally detrended for low frequency movement artefacts using Chronux’s locdetrend function with a 500 ms moving window and 100 ms overlap. Following local detrending, 50 Hz electrical mains noise was removed with Chronux’s rmlinesc function. LFP power spectrums (PSDs) and spectrograms were then calculated via the multi-taper spectral estimation method using the Chronux toolbox with a time-bandwidth product of 3 with 5 tapers, no FFT padding, and a bandpass filter of 0 to 100 Hz. The resulting power spectra covered 0 to 100 Hz in roughly 0.5 Hz increments. For a given recording session, all power spectrums and spectrograms were z-scored based on 100 randomly sampled 2 sec epochs from the pre-baseline period. Any trial where more than 15% of the frequency spectrum (0 to 48 Hz) measured over ± 3 standard deviations (SD) was removed. Likewise, trials were excluded if mean power measured over ± 3 SD within a frequency band of interest. While rare, these noisy trials tended to be caused by the rats hitting the headstage on the wall or arena floor, or from grooming behaviors prior to a pull attempt. Coherency, a measure of oscillatory synchronization, between the ACC and AI was analyzed using Chronux’s coherency function. Coherency was averaged across trials and data were pooled across sessions and rats. Spectrograms were created with Chornux’s mtspecgram function using a time-bandwidth product of 3, 5 tapers, a padding factor of −1 (no padding), and a 2 second moving window with 0.05 second step. One additional second of data was added before and after the 2 second epoch of interest to compute the spectrograms. The normalized LFP data from every attempt except for the first attempt within a weight block was averaged and then data were pooled across rats and sessions. We did not include the first attempt of a weight (e.g., trial 11, trial 21) as the rats were likely still expecting the previous weight prior to making an attempt. However, we can also not rule out with certainty that they were expecting the previous weight, e.g. they potentially registered the noises made by the experimenter while changing out the weight. Thus we have taken out the first attempt of every weight block (including 0 g) to control for the uncertainty in the rats’ expectations.

One-way ANOVA was used to test whether normalized power differed within a frequency band across rope weights with a significance value of 0.05. In addition to absolute weights, we also analyzed the data in terms of the rat’s relative performance state. For each individual session we assessed LFPs on: the start weight of 0g, the highest weight the rat achieved (i.e., completed 10 trials on), and the weight the rat quit on. One-way repeated measures ANOVA was used to test for changes in power and coherency across these three performance states within the session. Tukey’s Honestly Significant Difference Procedure was used to test for post-hoc multiple comparisons using Matlab’s multcompare function.

Our graphs attempt to show all the data points where feasible. However, we chose to programmatically limit the Y-axis based on the minimum and maximum quartiles of the data. As a result some individual data points outside of these ranges are not shown graphically. However, for coherency analyses, we hard coded the y-axis for all frequency bands for easier visual comparisons as coherency values can only range from 0 to 1. Color templates were made using ColorBrewer (https://colorbrewer2.org/).

This study was not preregistered. Data is not openly accessible but can be made available upon request.

### Code Accessibility

Analysis was carried out with custom MATLAB scripts. All code is available upon request to the corresponding author.

## Results

### Behavioral impact of progressive WLT

We first set out to characterize how rats behaved on the progressive WLT. Sixteen rats were tested on the progressive weight paradigm for a total of 114 sessions; each rat contributed 5-8 sessions. Within a session, rats generally completed the first three weights (0, 45, and 90 g) with ease before quitting on one of the latter three weights (135, 180 or 225 g; Figure 2a). The most common quit weight was 225 g, occurring in 33% of sessions. Rats’ average body weight was not correlated to the average weight they quit on (r = 0.02, p = 0.93). In 55% of sessions (51/114), rats completed at least one successful 30 cm pull on their quit weight before quitting, indicating that they were capable of successfully performing the action of lifting that weight up 30 cm but chose not to continue on. In 98% of sessions (112/114), rats were capable of at least lifting their quit weight off of the base. Together, rats’ pulling behavior indicated rats likely quit the task due to the increasing effort cost caused by the increasing weight. Furthermore, in 100% of sessions rats consumed the sucrose rewards during the end-of-session satiation check, suggesting that they did not quit the rope pulling task due to sucrose satiation.

**Figure 2:**
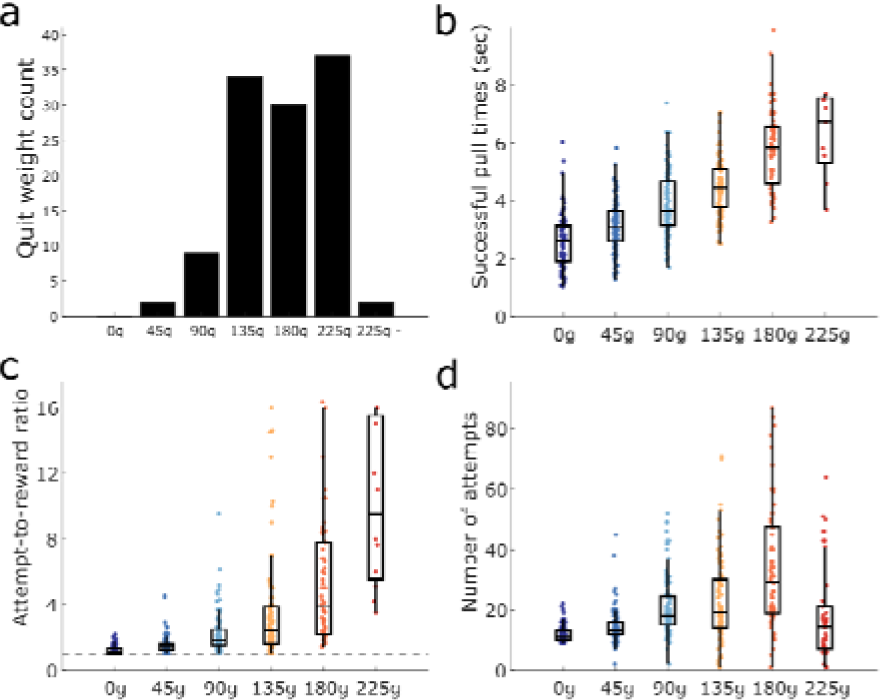
Progressive weight task behavior. a) Histogram of quit weight occurrence. “225g -” indicates the impossible phase where no rewards could be earned; see Methods. b) The time it took rats to pull the weights up 30 cm from the base, sufficient to trigger sucrose reward. Heavier weights took longer to lift. c) The attempt-to-reward ratio for each weight showing the average number of attempts rats made before completing a successful attempt. Dotted line represents a ratio of 1.0 where every attempt would be successful. Heavier weights took more pull attempts before a reward was successfully triggered. d) Average number of attempts rats made on each weight. Rats tended to make more attempts as weights got heavier, however, on heavier weights some rats made very few attempts before quitting.

The increasing effort load of the weights had a significant impact on the rats’ behavior. As the weight got progressively heavier, it took rats significantly longer to successfully pull the weight up 30 cm from the base (H (5, 469) = 232, p < 0.0001; Figure 2b). Pairwise comparisons between each weight and its preceding weight revealed significant increases in successful pull times across most weight pairs (p’s < 0.02), however, pull times between 0 g and 45 g (p = 0.06) and 180 g and 225 g (p = 1.0) were similar.

The increase in successful pull time with heavier weights indicates that the heavier weights are indeed more effortful for the rats. Successful pull times, however, can only be calculated for rewarded attempts and therefore do not fully capture the rats’ persistence at the task as it becomes increasingly difficult. In order to characterize the rats’ persistence we calculated an attempts-to-reward ratio of the average number of attempts made for each rewarded attempt for each weight. A ratio of 1.0 indicates every attempted pull was successful in triggering reward. Ratios greater than 1.0 indicate that the rats failed to pull the weight 30 cm on some attempts. Rats made significantly more attempts per reward as the weights increased (H (5, 482) = 268, p < 0.0001; Figure 2c). Pairwise comparisons revealed that rats generally made more attempts per reward as the weights got heavier (all p’s < 0.02), however, their performance was similar between 90 and 135g (p = 0.36) and 180 and 225g (p = 0.93). Thus, as the weights increase rats fail more often to successfully lift the weight.

On 69% of sessions where rats reached the 225 g weight they were not able to complete a single successful, rewarded pull (and thus we could not compute an attempt-to-reward ratio). To get a better idea of the rats’ persistence in the face of increasing effort demands, we measured the total number of attempts rats made on each weight. Weight had a significant impact on the number of attempts rats made (H (5, 543) = 169, p < 0.0001; Figure 2d). The average number of attempts increased weight-to-weight up to 180 g (p < 0.05), with the exception of 90 g to 135 g (p = 1.0), until 225 g where attempts significantly decreased on 225g compared to 180 g (p < 0.0001). Overall, these behavioral metrics indicate that the progressively increasing weights are more and more difficult for the rats to lift, requiring more effort (pull time, number of attempts) and persistence (attempt-to-reward ratio).

To further ensure rats were not quitting due finishing a specific number of pulls, sucrose satiation, or time-on-task, we ran 5 rats on a modified, fixed weight version of the WLT in the week after they ran the progressive weight paradigm. Rather than the weight progressively increasing, rats had 10 trials of 0 g followed by unlimited trials on 180 g. The 180 g weight was chosen as it was the maximal weight all 5 rats had completed 10 trials on in the progressive weight paradigm. Rats ran the fixed weight paradigm until they quit (no attempt for 2 minutes) or for one hour, whichever came first. Each rat completed 5 or 6 sessions of the fixed weight paradigm; 26 sessions were recorded in total. Compared to their progressive weight performance, during the fixed weight task the rats made significantly more attempts (median 202 attempts vs. 81 in the progressive weight; Z = −5.11, p < 0.0001; Figure 3a), earned significantly more rewards (68 vs. 42 in the progressive weight; Z = −4.73, p < 0.0001; Figure 3b), and performed the task for significantly longer before quitting (2123 sec vs. 1045 in the progressive weight; Z = −4.62, p < 0.0001; Figure 3c). Overall, these data further demonstrate that rats are quitting the progressive weight task due to the increase in effort (weights) and not due to repetitive rope pulling attempts, becoming satiated on sucrose, or quitting after an elapsed time.

**Figure 3:**
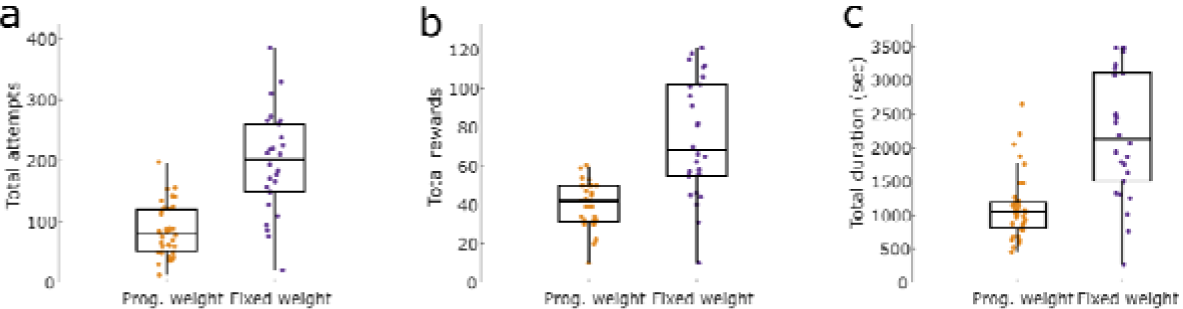
Behavioral comparisons between paradigms. In the progressive weight task, 45 g of weight was added to the rope after every 10 successful pulls. In the fixed weight task, a 180 g rope weight was accessible for up to 1 hour. In the fixed weight paradigm, rats made significantly more attempts (a), earned more reward (b), and performed for longer (c), as compared to their performance in the progressive weight paradigm.

### LFP correlates of progressive effort loading

We investigated the neural correlates of increasing effort costs by recording LFPs in the ACC and AI as rats carried out the progressive WLT. We specifically focused our analyses on the 2 sec prior to the rat lifting the weight up. This period reflects the rat choosing to actively engage in the task and make an attempt at expending energy to earn a reward. Changes in power in the delta (2-4 Hz), theta (7-9 Hz), beta (15-25 Hz), and gamma (55-100 Hz) frequency ranges were assessed. We predicted that low-frequency LFP power from the ACC would inversely correlate to effort costs, while low-frequency LFP power from the AI would positively correlate to effort costs.

The overall normalized ACC PSD for each weight is shown in Figure 4a. Weight had a significant influence on delta (F (5, 350) = 2.75, p = 0.019; Figure 4b) and theta band power (F (5, 350) = 2.97, p = 0.012; Figure 4c). Post-hoc multiple comparisons on delta and theta power using Tukey’s test revealed no significant pairwise differences (p’s > 0.05). Beta (F (5, 350) = 1.32, p = 0.257; Figure 4d) and gamma (F (5, 350) = 1.89, p = 0.095; Figure 4e) power were not significantly impacted by weight.

**Figure 4:**
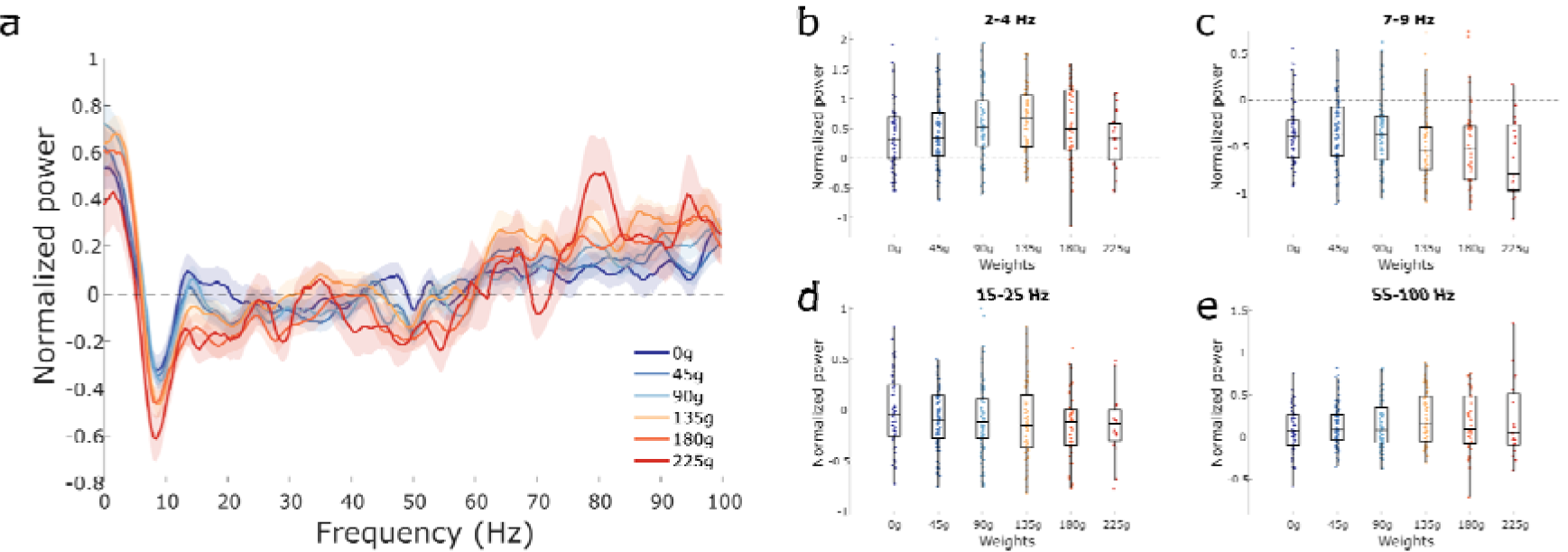
Changes in ACC LFP rhythms with increasing effort. a) Multi-taper power spectrum of ACC LFP across the six different weights. Shaded error bars represent ± 1 S.E.M. b-d) Normalized power across the four rhythms of interest (delta, theta, beta, and gamma, respectively). Box plots show first, second (median; middle line), and third quartiles. Whiskers are 1.5 times the interquartile range. All power calculations were normalized using the 2 min pre-baseline period, see Methods.

The overall normalized AI PSD for each weight is shown in Figure 5a. Weight had no effect on power within the delta (F (5, 352) = 0.97, p = 0.44; Figure 5b) or beta band (F (5, 352) = 1.23, p = 0.29; Figure 5d). Weight did have a significant effect on power within the theta (F (5, 352) = 2.38, p = 0.04; Figure 5c) and gamma bands (F (5, 352) = 4.80, p = 0.0003; Figure 5e). No post-hoc comparisons were significant for theta power (all p’s > 0.05). Post-hoc pairwise comparisons revealed significant differences between gamma power on 0 g compared to the other weights (p’s < 0.05), with the exception of 225 g (p = 0.054). No other pairwise comparisons were significant in the gamma band (p’s > 0.05).

**Figure 5:**
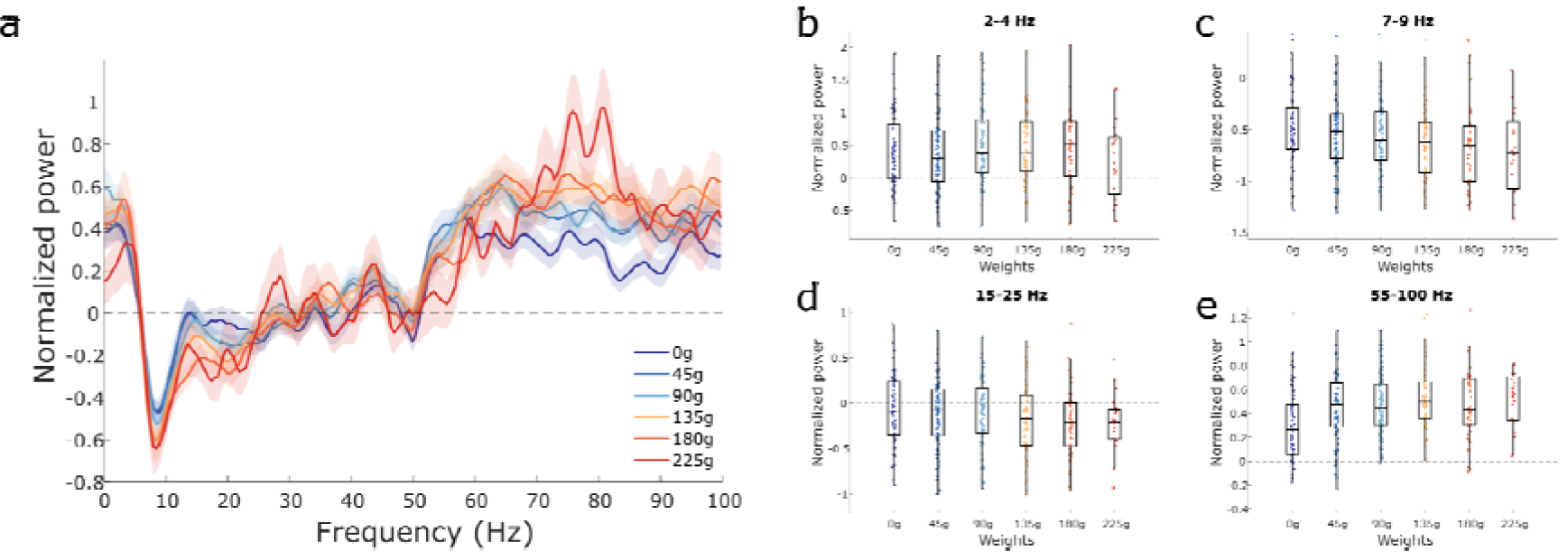
Changes in AI LFP rhythms with increasing effort. a) Multi-taper power spectrum of AI LFP across the six different weights. Shaded error bars represent ± 1 S.E.M. b-d) Normalized power across the four rhythms of interest (delta, theta, beta, and gamma, respectively). Box plots show first, second (median; middle line), and third quartiles. Whiskers are 1.5 times the interquartile range. All power calculations were normalized using the 2 min pre-baseline period.

### LFP correlates of quitting behavior

In addition to examining how increases in effort load (weight) affected regional activity in the ACC and AI, we were interested in examining how an animal’s performance state correlated to regional neural activity. As the WLT is a self-driven task, the animal can quit at any point, on any weight. This generates inter- and intra-individual variability in quit points, and quit point metrics, likely reflecting variability in motivational states. Due to the rats’ quitting behaviors, the motivational state of a rat on a given weight could be different to that of another rat on the same weight. For example, one rat may quickly complete 10 successful pulls on the 135 g weight, while another rat may quit on 135 g after only a few attempts. To address session-to-session variability, we divided up the rats’ performance based on their relative achievement for a given session. For each individual session we assessed LFPs from three blocks of interest: 1)We analyzed the specific frequency bands during the low-effort start weight of 0 g; 2) the rat’s achievement weight (i.e., the maximum weight that the rat completed 10 trials on); and 3) the weight the rat quit on.

The decision to quit is likely a dynamic process, occurring as animals assess their current internal states with the costs and benefits of the currently available task. We were interested in visualizing the LFP temporal dynamics as rats chose to carry out their attempts across the three performance states (0 g, achievement weight, and quit weight). Figure 6a shows the averaged normalized spectrograms across all recordings of the three performance states. Prominent increases in theta and gamma power can be seen in the achievement and quit weight compared to the 0 g weight. In contrast, theta appears to decrease in power across performance states. The overall normalized ACC PSD across the three states can be seen in Figure 6b. When each frequency band of interest was independently assessed, significant differences in power were observed between performance states in the delta rhythm (F (1.96, 95.9) = 16.8, p < 0.0001; Figure 6c), theta rhythm (F (1.64, 80.1) = 5.45, p = 0.01; Figure 6d), beta (F (1.58, 77.3) = 4.22, p = 0.026; Figure 6e), and gamma rhythm (F (1.53, 74.8) = 11.49, p = 0.0002; Figure 6f). Post-hoc comparisons revealed a significant increases in ACC delta and gamma power between 0 g and the achievement weight (p < 0.0001 and p = 0.0002, respectively). Both theta and beta power significantly decreased from the achievement weight to the quit weight (p = 0.03 and p = 0.02). Every brain rhythm showed a significant difference from 0 g to the quit weight where theta (p = 0.02) and beta (p = 0.03) power decreased and delta (p = 0.0003) and gamma (p = 0.004) power increased.

**Figure 6:**
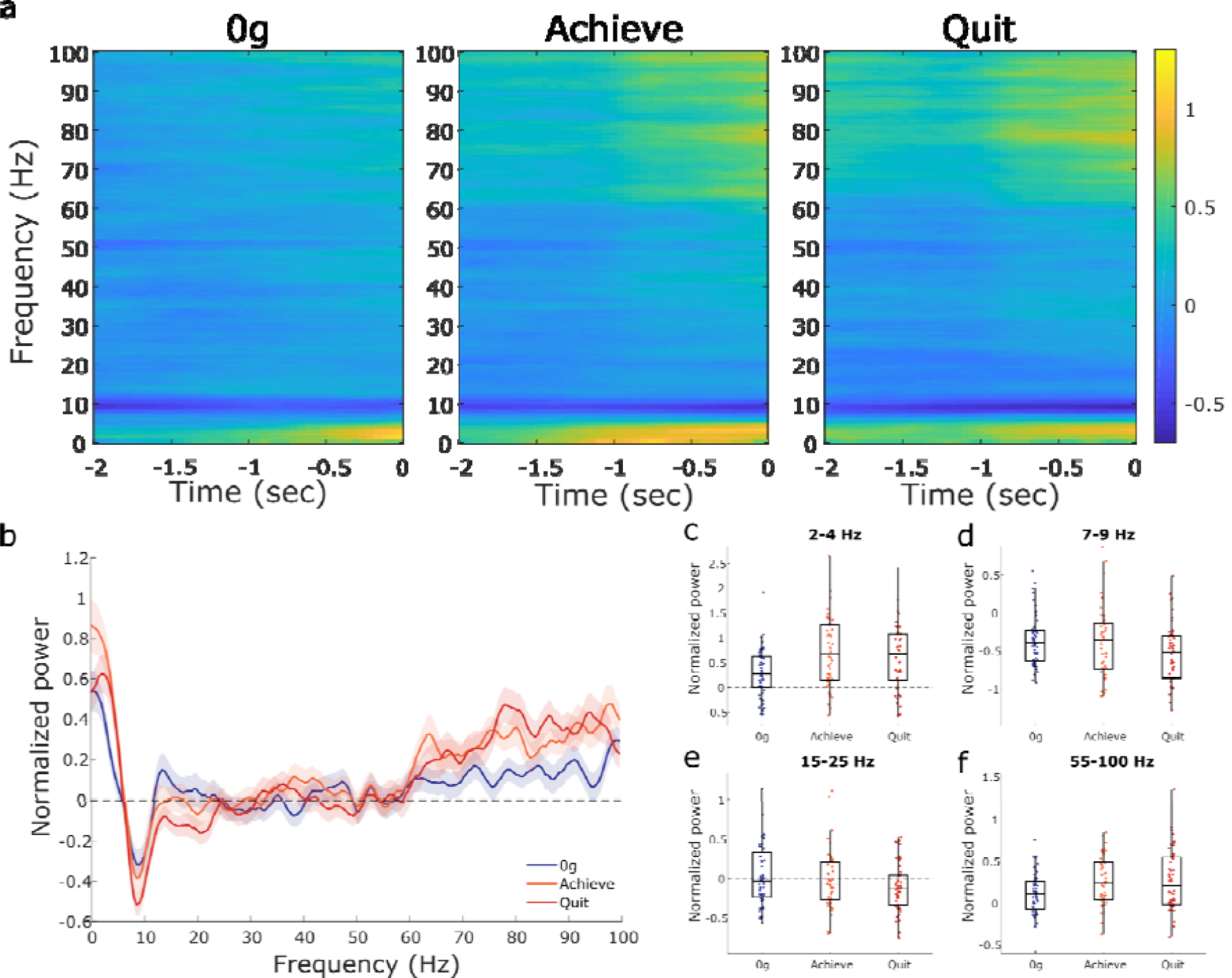
Changes in ACC LFP rhythms across performance states. a) Average ACC spectrograms for the three performance states showing mean normalized power in the 2 seconds prior to the attempt. 0 g (blue; b-f) is the 0 g weight. The Achieve weight (orange; b-f) is the last complete (10 successful pulls) weight while Quit (red; b-f) is the weight the rats quit on. b) Multi-taper power spectrum of ACC LFP across the three performance states. Shaded error bars are ± 1 S.E.M. c-f) Normalized power across the four rhythms of interest (delta, theta, beta, and gamma, respectively). Box plots show first, second (median; middle line), and third quartiles. Whiskers are 1.5 times the interquartile range. All power calculations were normalized using the pre-baseline period.

Figure 7a shows the average normalized AI spectrograms from 0g, achievement weight, and quit weight. Clear changes from the LFP baseline are apparent at the delta, theta, and gamma rhythms. These shifts in power seem to be most prominent just prior to the rat pulling the rope. The overall normalized AI PSD across performance states is shown in Figure 7b. Power in the delta frequency range (F (1.73, 84.8) = 2.1, p = 0.14; Figure 7c) did not change significantly across the three performance states. In contrast, AI theta (F (1.73, 84.8) = 17.5, p < 0.0001; Figure 7d), beta (F (1.73, 84.8) = 5.55, p = 0.0077; Figure 7e), and gamma rhythms (F (1.74, 85.4) = 19.8, p < 0.0001; Figure 7f) did show significant differences across performance states. Post-hoc pairwise tests revealed a significant decrease in theta power from 0 g to the achieve weight (p = 0.0007) and quit weight (p < 0.0001) as well as from the achieve weight to the quit weight (p = 0.042). Beta power decreased significantly only from 0 g to the quit weight (p = 0.009). In contrast, gamma power increased from both 0 g to the achievement weight (p < 0.0001) and from 0 g to the quit weight (p < 0.0001).

**Figure 7:**
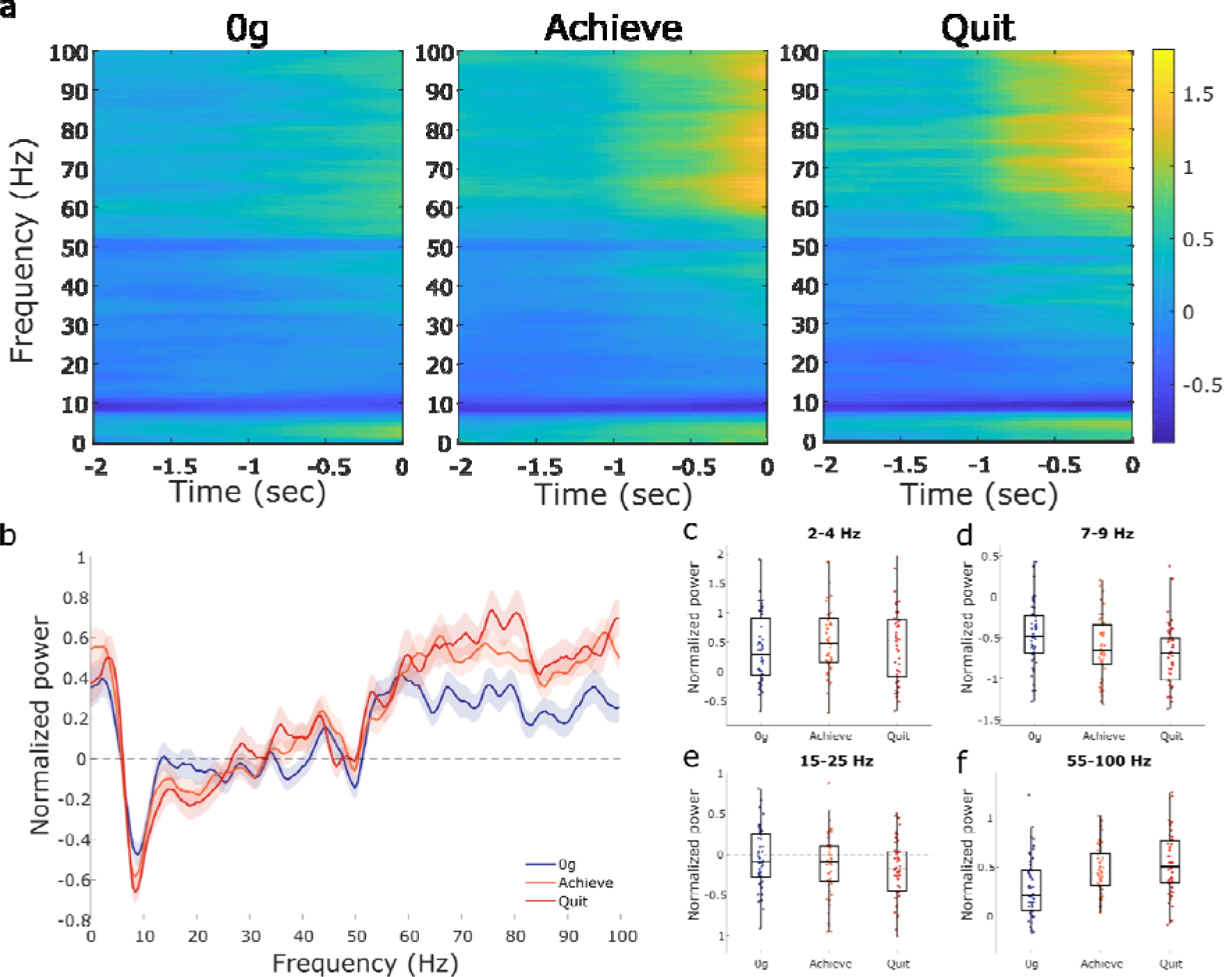
Changes in AI LFP rhythms across performance states. All power in normalized based on the pre-baseline period. a) Average AI spectrograms for the three performance states showing mean normalized power in the 2 seconds prior to the attempt. 0 g (blue; b-f) is the 0 g weight. The Achieve weight (orange; b-f) is the last complete (10 successful pulls) weight while Quit (red; b-f) is the weight the rats quit on. b) Multi-taper power spectrum of AI LFP across the three motivational states. Shaded error bars are ± 1 S.E.M. c-f) Normalized power across the four rhythms of interest (delta, theta, beta, and gamma, respectively). Box plots show first, second (median; middle line), and third quartiles. Whiskers are 1.5 times the interquartile range.

### ACC-AI coherency shifts across performance states

The activity of distant brain regions can become synchronous during specific behaviors, possibly reflecting task-specific information processing in both regions (Harris and Gordon 2015). We were interested in whether or not ACC-AI coherency increased as the animal got closer to quitting, with the ACC evaluating effort costs in the context of AI’s interoceptive information. To determine if ACC-AI coherency changes occurred across WLT performance states, we assessed coherency during the two second attempt epoch across our three states of interest: 0 g, achievement weight, and quit weight. Coherency in the delta range showed a significant effect across states (F (1.82, 27.4) = 11.38, p = 0.0004; Figure 8a). Post-hoc showed coherence significantly increased from 0 g to the achievement weight (p = 0.03) and quit weight (p = 0.002). There were no significant changes in ACC-AI coherency in the theta (p = 0.67; Figure 8b), beta (p = 0.38; Figure 8c), or gamma (p = 0.10; Figure 8d) rhythms when performance states were compared.

**Figure 8:**
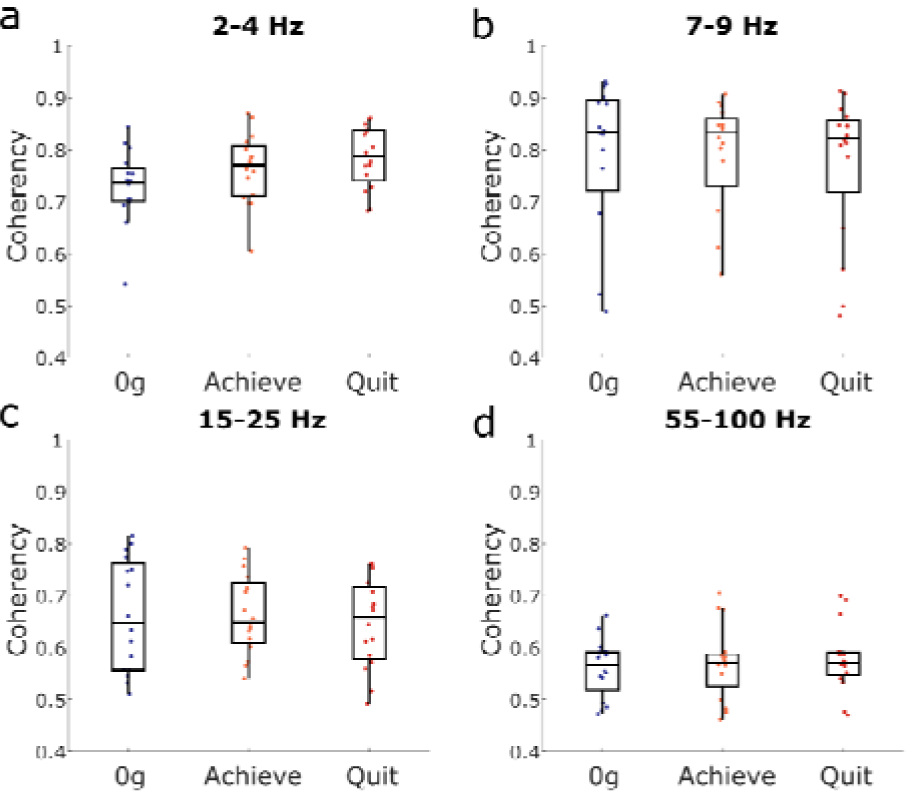
ACC-AI coherency measures. Coherency changes across performance states in the delta (a), theta (b), beta (c), and gamma (d) rhythms. Box plots show first, second (median; middle line), and third quartiles. Whiskers are 1.5 times the interquartile range.

## Discussion

Behaviorally, this study demonstrates the efficacy of our novel WLT for probing effort exertion in laboratory rats. Progressively increasing the weight the rats must lift in order to earn a fixed reward resulted in distinct behavioral shifts, reflective of increasing effort cost (see Figure 2). Heavier weights (≥ 135 g) proved to be particularly effortful, and resulted in high rates of quitting. As reward amount was held constant, the observed quitting behaviors were most likely due to a decrease in perceived task utility driven by the increased physical effort costs. Time expenditure in the task was not likely to account for the quitting behaviors we observed, as experiments using a fixed-weight paradigm demonstrated that some rats will rope pull 180 g for up to 1 hour (Figure 3c). Motivated sucrose intake at the end of the session (sucrose satiation check; see Methods) and rats consuming significant more sucrose on the fixed weight paradigm (Figure 3b) further support the interpretation that quitting the progressive WLT was associated with mounting task costs, rather than reward devaluation due to satiation.

Only male rats were used in this study. However, previous research has demonstrated sex differences in decision making behaviors (Orsini and Setlow 2017). For example, Orsini et al. (2016) utilized a risky decision making task and found that male rats were more risk seeking than female rats. While no direct sex difference comparisons have been conducted on effort discounting in rats, Uban et al. (2012) demonstrated estrogen receptor agonists shift behavioral preferences away from high-cost, high-reward towards low-cost, low reward choices. As our present study only used male rats, our results may not generalize to female rats on the WLT.

Neurophysiologically, we predicted that low-frequency LFP power from the ACC would inversely correlate to effort costs, while low-frequency LFP power from the AI would positively correlate to effort costs. When the progressive WLT was examined based on absolute weight categories, our hypotheses were only partially supported. In the ACC, both delta and theta power had a main effect of weight but no post-hoc tests were significant. Thus, we cannot conclude whether or not increasing effort costs increase low frequency rhythms in the ACC. In contrast, AI LFP power did not show a progressive increase in any of the low-frequency bands examined as we had predicted. There was a suspected decrement in theta power in response to rope weight upon first visual inspection (see Figure 5a), but this was not significant. Regional gamma power in the AI did significantly increase from 0 g to the heavier rope weights, notably in the 75-85 Hz range.

While we found changes in ACC delta and theta along with AI gamma across the six weight conditions of the WLT session, one shortcoming of analyzing the six weight conditions as a single factor is that there is considerable behavioral variability occurring, notably on the higher weights. For example, 135 g may be a ‘mid-way’ weight for Rat A who proceeds all the way to the 225 g in a given session, but 135 g may be the ‘quit point’ for Rat B in a given session. Accordingly the neural states would likely be very different between Rat A and Rat B in those sessions, despite the 135 g condition being nominally the same. For this reason we performed a secondary analysis on the dataset on the basis of performance state rather than specific weight.

Performance state analysis demonstrated a wider range of impacts across multiple neural rhythms. In the ACC, delta power was lower during the initial 0 g weight compared to the achievement and quit weight, and there was an increase in delta rhythm coherency between the ACC and AI across performance states. In general, slower rhythms are thought to reflect network level coordination across many different brain regions (von Stein and Sarnthein 2000; Buzsaki and Draguhn 2004). The increase in coherency on the challenging weights (achieve and quit) compared to the 0 g weight may reflect the increased network activity within the salience network as the WLT becomes increasingly demanding (Uddin 2015). Furthermore, delta rhythms have been shown to coordinate prefrontal regions with VTA dopamine circuitry (Fujisawa and Buzsaki 2011; Elston and Bilkey 2017). Dopamine may help drive continued task performance in the WLT in the face of increasing effort conditions by biasing ACC networks to discount effort costs and/or amplify reward benefits (Schweimer and Hauber 2006).

In both the ACC and AI, theta and gamma power showed significant but opposing changes in relation to performance state: regional theta power decreased and gamma power increased as the animal progressed towards their quitting point for that session (see Figures 6, 7). In both the ACC and AI, theta powered dropped significantly on the quit weight compared to 0 g and the achievement weight. The decrease in theta power on the quit weight may reflect the shift in task utility and the task no longer being “worth-it” compared to the 0 g and achievement weight where utility is high. In contrast, both brain regions’ gamma power showed a significant increase from task onset (0 g) to achievement weight and quit weight. Thus, gamma appears to become elevated earlier on in the task (achievement weight) and stay elevated relative to 0 g. In general, the elevated gamma power may reflect the increased local neuronal activity as the task becomes more challenging. Overall, these shifts in theta and gamma power may reflect changing motivational states as the rat approaches the decision to quit the task.

Previous studies investigating the neural correlates of effortful or persistence-like behaviors have tended to run subjects for a fixed time or for a fixed number of trials (Croxson et al. 2009; Hillman and Bilkey 2012; Engstrom et al. 2014; McGuire and Kable 2015; Hart et al. 2017). However, these studies do not capture what occurs in the brain when subjects voluntarily quit a task. During physically demanding tasks, ACC single unit population activity in rodents and non-human primates has been shown to reflect the current value of a given behavior (Kennerley et al. 2009; Cowen et al. 2012; Porter et al. 2019). Our LFP data are in line with these findings, showing that in the rat ACC and AI, theta power decreases from the start of the task (0 g) to the self-selected end of the task (achievement weight and quit weight). Theta rhythm may reflect an overall net utility of the action at hand, and relatedly, motivation to continue performing the task or to quit. Elston and Bilkey (2017) found a similar result in rats using a fixed reward, jumpable-barrier effort task: ACC theta power increased on low-effort (no barrier) trials, and decreased on high-effort trials (barrier present). Overall, ACC theta power may reflect the current value of the behavior being carried out and the cognitive control exerted by the ACC to bias behavior towards particular actions (Womelsdorf et al. 2010).

These rat-based observations and our interpretation as to why theta power decreases across the progressive WLT may initially appear at odds with human data. In EEG studies, frontal midline theta reliably increases as task difficulty and/or duration increases (Paus et al. 1997; Smit et al. 2005; Mitchell et al. 2008). One possible concordant interpretation links back to the notion of running subjects for a fixed duration versus allowing subjects to self-select the endpoint. If theta power in the ACC is reflective of utility of a current action, theta power progressively decreases in our rat task as the task gets difficult yet the reward amount remains the same; utility progressively declines. Human subjects, however, have additional factors feeding into cost-benefit utility calculations and these include completion goals. Social contract pressure to not ‘give up’ or quit during a research task of fixed duration – particularly if under researcher supervision – would increase the utility of performing and persisting in difficult tasks. Likewise nearing the end of a session of known duration may confer lowered cost, and/or added benefit to task trials, lending heightened utility to trials proximal to the end point. Such factors may in part account for why midline frontal theta increases in human EEG studies, and would be an interesting area for future investigation.

In contrast to the decrement in theta power across performance states, power in the gamma band in the ACC and AI increases across performance states. Cortical gamma has been proposed to be generated by interneurons and the need to balance excitation and inhibition during periods of activation (Bartos et al. 2007; Merker 2016). Increased gamma power in our WLT may reflect the increasing cognitive processing demands of the task; mountings costs must be dynamically integrated with interoceptive signals, such as hunger and fatigue, to compute on-going utility. For example, gamma rhythms can selectively amplify or dampen incoming signals dependent on when in the gamma cycle the information arrives (Fries et al. 2001; Sohal et al. 2009). Such a mechanism could reflect the increased information processing needed as the task becomes more challenging. Indeed, gamma power in the ACC and AI shows an increase in power during the achievement and quit weights compared to the easiest, and highest utility, 0 g weight. This may be reflective of the AI’s role in integrating the physical cost of behaviors, and in particular the negative affective aspects of those physical costs, with the dynamic changes in internal states as the rats approach their quitting point (Williamson et al. 1999; Prevost et al. 2010).

Similarly, ACC’s heightened gamma power alongside increasing effort costs may be reflective of its role in effort-based cost-benefit decision making (Kennerley et al. 2006; Croxson et al. 2009; Klein-Flugge et al. 2016) and cognitive-control (Lorist et al. 2005; Woodward et al. 2008; Holroyd and McClure 2015). For example, foraging tasks in humans (Kolling et al. 2012) and non-human primates (Hayden et al. 2011) demonstrate increasing ACC activity as subjects approach the decision to search for another foraging patch. While subjects in these foraging-based tasks do not quit the testing session, these data provide evidence that the ACC may determine the value of “quitting” the current behavioral option. Since the gamma rhythm is thought to reflect local circuit activity and gate incoming information (Fries et al. 2001; Sohal et al. 2009), the shifts in gamma activity across the task may reflect the ongoing integration of many sources of information (e.g. fatigue, satiation, reward history, current effort cost) to determine whether or not the task is worth persisting at or quitting.

Our WLT demonstrates that the rat’s behavior is altered as effort costs increase while reward stays constant. As shown in Figure 2, rats struggle to lift heavier weights, fail more often on their attempts, and, as result of failing more often, make more attempts as the weight increases. These factors likely contribute to the rat’s choice to quit and may also contribute to the changes in LFP power we report here. For example, due to the rats failing more often, they earn rewards at a slower rate. These failed attempts may also mean the rat perceives the task as becoming more volatile as their attempts become less likely to result in a reward. Both reward history (Kolling et al. 2016) and task volatility (Behrens et al. 2007) have been shown to be represented by the ACC.

Both ACC and AI activity changes may likewise be altered by the increasingly infrequent sucrose consumption (Horst and Laubach 2013; Becker et al. 2017). In particular, a proposed role of the beta rhythm is in the maintenance of the status quo in that the current sensorimotor plan is expected to be maintained (Engel and Fries 2010). However, as the task becomes more difficult, rewards more infrequent, and fatigue sets in, the status quo of carrying out the WLT may be challenged. Indeed we observed a decrease in beta power on the quit weight compared to 0 g in both brain regions. Thus, factors associated with the changes in weight, changes in behavior, and changes in task outcome may be contributing to the observed shifts in ACC and AI LFPs.

Here was have characterized LFP activity in the ACC and AI during a physical effort-based task, where rats have the choice to engage in the task or quit the session. Motivational performance state, rather than absolute effort cost (weight), appears to better capture the dynamic changes in ACC and AI LFP power observed across the session. As rats approach the decision to quit, ACC and AI theta power decreases while gamma power increases. Our results may guide future effort-based cost-benefit decision studies in targeting the source and content of these rhythms.

## Acknowledgments

This study was supported by Marsden Fund grant U001617 (K.L.H.) from the Royal Society of New Zealand Te Apārangi.

